# Genotype-by-diet interactions determine Black Soldier Fly life-history traits

**DOI:** 10.64898/2026.04.21.719825

**Authors:** Jessica Jiogue-Lacdo, Marie Merle, Mhant Kondé, Meroua Foughar, Chloé Genevey, Agus D. Permana, Pierre-Olivier Maquart, Jonathan Filée

## Abstract

The black soldier fly, *Hermetia illucens*, is increasingly valued in applied entomology due to its remarkable capacity to upcycle organic waste and for high nutritional value of its larvae. As a result of global expansion and domestication, the species now displays substantial genetic diversity, yet performance differences between strains remain poorly documented. This study aimed to better understand the relationship between genotype and phenotype, as well as their interaction, to support the improvement of its domestication. Five distinct strains collected from the wild by artisanal farmers or obtained from industrial farms were genetically characterized using whole genome sequencing. These analyses revealed high genetic divergence based on mitochondrial genome and SNP nuclear genome phylogeny. To assess phenotypic performance, the strains were reared on three diets differing in nutritional value: poor (alfalfa meal), intermediate (wheat bran) and rich (chicken feed) and their growth rate was assessed. At harvest, we evaluated different life history traits including survival rate, average larval mass, feed conversion ratio, substrate reduction and bioconversion rate. Statistical analyses revealed strong effects of both diet and strain (p < 0.001), but the key result was the pronounced strain × diet interaction. Performance varied drastically depending on substrate quality: some strains showed high versatility across all diets, while others performed mainly on nutrient-rich substrates or excelled in substrate degradation. In contrast, other strains displayed more specialized profiles, with marked sensitivity to fibrous diets. These contrasted reaction norms highlight that diet performance cannot be interpreted independently of the strain genetics. Overall, these findings underscore the value of preserving diverse local genetic resources and the need for improved molecular tools to guide strain selection.

**Implication:** This study shows that performance of the black soldier fly depends strongly on interactions between genetic background and diet, confirming the importance of genotype–environment relationships. While results are based on a limited number of strains and substrates, the consistent strain × diet interaction suggests broader relevance for rearing systems. These findings highlight the need to integrate genomic data into phenotypic assessments. Practically, they indicate that strain selection should be tailored to substrate type to optimize productivity and efficiency. This has direct economic benefits for insect farming and waste management industries because improved strain–diet matching can enhance organic waste bioconversion and support circular economy strategies. Overall, preserving genetic diversity and developing molecular tools for strain selection are key steps toward more sustainable and efficient insect production systems of this study have implications for the development and sustainable BSF systems production.

## Introduction

The global human population is expected to exceed 10 billion by 2050, with more than half of this growth occurring in Africa (United Nations, 2015). Meeting the resulting food demand will require a substantial increase in agricultural productivity. According to the Food and Agriculture Organization (FAO), global food production must increase by approximately 70% to ensure food security, a challenge further compounded by rising income levels and associated shifts in dietary preferences toward animal-derived protein sources (FAO, 2009; van Huis *et al*., 2013). Consequently, the demand for animal feed protein is projected to rise sharply, necessitating the development of sustainable and resilient feed production systems. Currently, animal feed formulations rely heavily on fishmeal and soybean meals, both of which present significant economic and environmental limitations. Fishmeal production intensifies pressure on marine ecosystems through overfishing, while soybean cultivation is a major driver of deforestation, biodiversity loss, and greenhouse gas emissions, particularly in South America (Tacon & Metian, 2008; Rana *et al*., 2015). In addition, both feed ingredients are characterized by high price volatility and limited accessibility for small-scale producers. In many low- and middle-income countries, dependence on imported feed ingredients further exacerbates vulnerability to global supply disruptions and economic instability (van Huis *et al*., 2013).

In this context, insects have emerged as a promising alternative source of animal protein for feed applications. Among them, the Black Soldier Fly (BSF), *Hermetia illucens* (Linnaeus, 1758), has attracted particular attention due to its high feed conversion efficiency, ability to valorize organic waste, and suitability for large-scale production (Cickova *et al*., 2015; Pastor et *al*., 2015). BSF larvae are increasingly recognized as highly digestible and nutritionally valuable ingredients for aquaculture, poultry, and pig diets (Oonincx *et al*., 2015; Barragan-Fonseca *et al*., 2017). However, rearing BSF on different diets leads to substantial variation in larval performance and nutritional composition suggesting that rearing conditions and biological factors strongly influence zootechnical outcomes (Trinh *et al*., 2015). While performance variation in relation with diet and rearing conditions have been extensively studied in the BSF, the role of the genetic background received much less attention. The comparison of three different strains (Wuhan, Ghanzou and Texas) indicate important variation of relevant life history traits as development speed or waste conversion ratio (Fen Zhou et al., 2013) Similarly, comparison of three industrial strains reveals strong phenotypic variation in terms of body size suggesting strong genotype by diet interaction (Generalovic *et al*., 2025). If the two latter studies lack genetic characterization of different strains, the work of Sandrock *et al*. was the first to combine genetic characterization using microsatellite genotyping of 4 strains from diverse geographical origins and estimation of commercially important life history traits (Sandrock *et al*. (2022). This study reported pronounced strain-specific responses to nutrient-rich substrates, whereas performance converged under low-quality dietary conditions. It also demonstrates that a strain is not universally better than the other because each strain appears best suited to specific conditions, highlighting the need for a better understanding of the genetic architecture of each trait.

On a broader scale, genetic studies have revealed substantial global structuring in BSF populations, with strong geographic differentiation and deep evolutionary divergence among lineages (Stahls *et al*., 2020; Kaya *et al*., 2021; Guilliet *et al*., 2021, Silvaraju *et al*., 2026). Experimental evidence has demonstrated that mass rearing can reduce genetic diversity and lead to rapid divergence between wild and captive populations (Hoffmann *et al*., 2021). In addition, the domestication process induced dramatic reduction of the genetic diversity in relation with population bottleneck in addition to balancing selection and selective sweeps processes. (Silvaraju *et al*., 2026). Similarly, Gligorescu *et al*. (2023) demonstrated rapid adaptive responses of BSF populations to low-quality diets over multiple generations, highlighting trade-offs between adaptation and overall performance. Despite this growing body of genetic knowledge, the extent to which genetic differentiation interacts with diet quality to shape zootechnical performance remains poorly understood.

In this study, we address this gap by systematically assessing genotype-by-environment interactions in BSF using a fully crossed factorial design. Five genetically distinct BSF strains were reared on three nutritionally contrasting diets to evaluate strain-specific responses and their interactions across key larval performance traits, including survival, feed conversion, substrate reduction, and bioconversion efficiency. By integrating genetic diversity with dietary context, this work aims to provide new insights into how genetic background influences larval performance under different nutritional environments. These results may contribute to improving strain selection and optimizing rearing strategies for sustainable industrial BSF production.

## Materials and methods

### Strains origins

BSF larvae used in this study originated from two distinct breeding systems: artisanal and industrial stocks (Figure 1). The artisanal breeding stock included three strains originating from small-scale production systems: (i) a strain from Myanmar collected from an artisanal farm in Rangoon region, (ii) a strain from Indonesia obtained from a small private breeding facility in Java, and (iii) a strain from Cameroon collected from a small-scale production unit located in the Dschang region. The industrial breeding stock included two strains originating from large-scale commercial production systems: (i) a strain from China commonly referred to as the “Wuhan” strain, widely used in industrial BSF production in Asia, and (ii) an European strain, extensively used in large-scale commercial farms across Europe and the United States. All the strains were reared in a colony maintained separately prior to the experiment to preserve their genetic identity.

**Figure 1:**
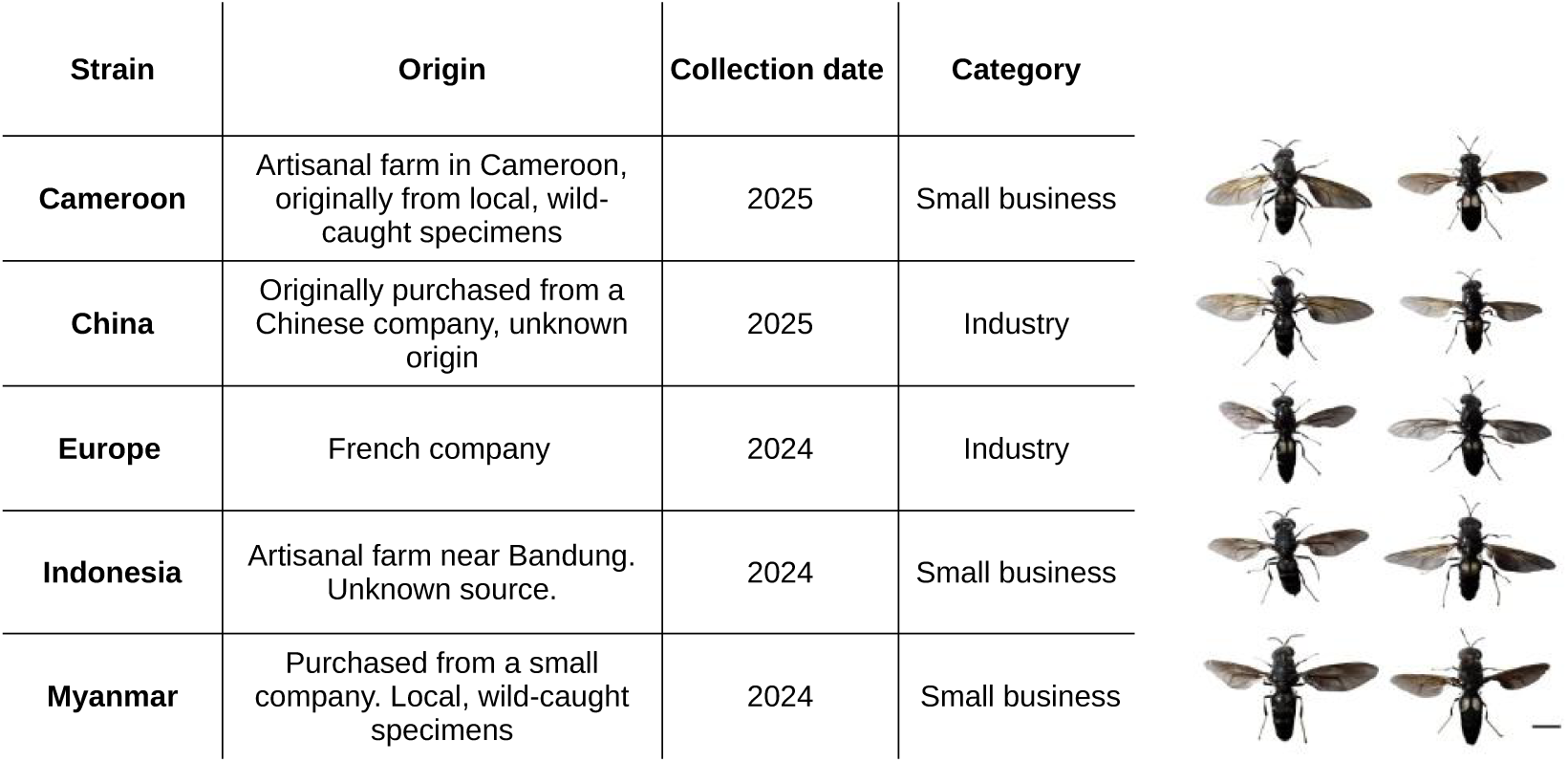
Main characteristics of the five BSF strains used in this study. Photographs show one male and one female reared under identical laboratory conditions. Scale bar: 5 mm.

### DNA extraction

Genomic DNA was extracted from adult BSF using the NucleoMag Tissue DNA kit (Macherey-Nagel) according to the manufacturer’s protocol. For each specimen, wings and abdomen were removed prior to extraction to facilitate tissue homogenization and avoid potential contamination from eggs. Samples were finely homogenized and incubated in 200 µL of lysis buffer (T1) with 25 µL of Proteinase K at 56 °C for 1–3 h. Following centrifugation, RNase was added. DNA purification was then performed using magnetic beads on a MagnetaPure 32 Plus automated extraction system, including successive washing steps (MB3, MB4 and MB5 buffers) and a final elution in 50 µL of the elution buffer (MB6). DNA quantity and purity were assessed using Nanodrop spectrophotometry and Qubit fluorometric quantification.

### Mitochondrial genome phylogeny

Mitochondrial genome sequences were assembled from Illumina paired-end sequencing reads using GetOrganelle (v1.7.7.1). The resulting circular mitochondrial genomes were linearized by sequence rotation prior to alignment. These sequences were combined with 60 complete *Hermetia illucens* mitochondrial genomes retrieved from Guilliet *et al*. (2022). Multiple sequence alignment was performed using MAFFT aligner included as a plug-in in the Geneious software. We used the algorithm G-INS-i with a scoring matrix of 200 PAM/k = 2, a gap open penalty at 1.53, and an offset value at 0.123. We use the MEGAX (Kumar et al. 2018) software to determine the best model and compute a maximum likelihood tree using IQ-TREE (Minh *et al*., 2020) with 1,000 Ultrafast Bootstraps Approximation (Minh et al., 2013). Phylogenetic inference was conducted The resulting phylogenetic tree was visualized and annotated using the Interactive Tree of Life: iTOL (Letunic.2021). The tree was rooted at midpoint.

### Variant calling and whole-genome nuclear SNP phylogeny

Raw Illumina sequencing reads were aligned to the BSF reference genome (NCBI accession: GCF_905115235.1) retaining only the nuclear chromosomes using minimap2 (version 2.30) (Li *et al*., 2018). Alignment files were converted to BAM format, sorted, and indexed using SAMtools (Li *et al*., 2009). Variant calling was performed using the bcftools mpileup pipeline. Raw variants were filtered using vcfutils.pl varFilter with a minimum variant quality threshold of QUAL ≥ 20 and a read depth ranging between 10 and 50 (−d 10, −D 50), retaining high-confidence biallelic SNPs.

For nuclear phylogenetic reconstruction, filtered SNPs were converted to PHYLIP format using vcf2phylip.py, retaining only variable sites. Maximum likelihood phylogenetic inference was conducted using IQ-TREE (Minh *et al*., 2020), under the GTR+ASC substitution model, according to BIC, to account for ascertainment bias associated with variable-site-only alignments. Branch support was assessed using 1,000 ultrafast bootstrap replicates. In parallel, the level of genomic similarity was performed based on common shared SNPs between strains using the distance matrix and the nuclear genetic structure was explored using principal component analysis (PCA) using VCF2PCACluster (v2.8) (Zhang *et al*., 2023), and results were visualized in three-dimensional space.

### Experimental diets and rearing conditions

Three experimental diets were used to rear BSF larvae: chicken feed (CF), alfalfa meal (AM), and wheat bran (WB). These diets were selected to represent distinct nutritional levels based on previous experiments conducted in the laboratory. The complex, high-protein diet consisted of 100% chicken feed (CF; 17% crude protein). The intermediate and low-protein diet consisted of single source alfalfa meal (AM; 16% crude protein) and 100% wheat bran (WB; 12% crude protein) respectively. The detailed composition of the experimental diets is provided in Table 1. For each strain and each diet, five independent replicates were established, each consisting of 200 larvae and 500g of substrate. Larvae were reared on the respective substrates under controlled environmental conditions at 27.0 ± 0.5 °C and 70 ± 5% relative humidity in a climate-controlled insectarium.

**Table 1:** Composition of different diets.

| Composition | Chicken feed (%) | Wheat bran (%) | Alfalfa meal (%) |
| --- | --- | --- | --- |
| Dry matter | 9,17 | 16,16 | 8,6 |
| Ash | 0,946 | 0,965 | 0,887 |
| Proteins | 17,16 | 12,03 | 16,23 |
| Lipids | 4,33 | 6,59 | 6,86 |
| Carbohydrates | 68,39 | 64,25 | 67,42 |

### Larval growth and performance measurement

Larval growths were monitored using batches of 200 five-day-old larvae reared in plastic containers (20 x 30 x 15cm) under control conditions. For each replicate, 20 larvae were randomly sampled every two days throughout larval development. Sampled larvae were rinsed with lukewarm tap water to remove residual substrate, wrung and weighed. Larval weight measurements were continued until 25% of individuals within a replicate had reached the pre-pupal stage. The day at which larvae attained their maximum mean body weight was defined as the end of the larval feeding stage. Following this point, larvae entered the pre-pupal stage and stopped feeding, at which time the experiment was terminated.

The collected data was used to quantify larval growth dynamics and developmental timing traits for each strain - diet combination.

Survival rate of larvae, representing the proportion of live larvae at a given time compared with the initial number of larvae :

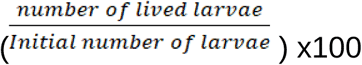

The average daily gain, this parameter made it possible to assess the growth rate of the larvae throughout their rearing period and was calculated by:

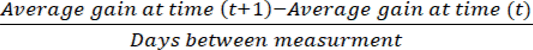

Feed Conversion Ratio appears which allows us to know if the larvae have properly assimilated the food that was given to them. The lower it is, the more productive the larvae are. It were calculated taking account humidity on both larvae and feed according to Diener *et al*. (2009), using the formula:

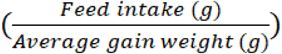

The reduction of the substrate is a parameter that allows measuring the effectiveness of the given food by determining the proportion consumed compared to the initial dry matter percentage. The higher it is, the more the larvae will have valued the substrate. According to Diener *et al*., (2009), it is calculated with the formula:

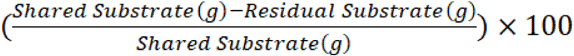

The bioconversion rate allows measuring the proportion of dry matter from the distributed substrate that has been transformed into fresh larval biomass. The higher it is, the more effective the conversion was. It is calculated according to Ebeneezar *et al*., (2021), it is obtained by:

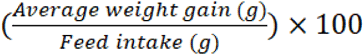

The emergency rate of adults’ flies were evaluated by isolating 50 random pupae per batch (four replicates); from the end of growth phase until the flies emerge.

### Statistical analysis

All statistical analyses and data visualizations were performed in R (v4.4.2; https://www.r-project.org/). Diet effects within each strain were assessed by one-way ANOVA, and strain effects, diet effects, and their interaction were evaluated jointly by two-way ANOVA. Normality and homoscedasticity of residuals were verified by Shapiro-Wilk and Levene tests, respectively. When a main effect or interaction was significant (α = 0.05), Tukey’s honestly significant difference (HSD) post hoc test was applied to identify pairwise differences between means.

## Results

### Strain phylogeny and genetic diversity

Albeit originating from 3 different continents, the five BSF strains used in this study look very similar with no clear differentiation regarding the body shape, coloration or wing patterns (Figure 1). To characterize the genetic diversity among these strains, we first build a maximum likelihood phylogenetic tree using complete mitochondrial genome sequences of the five strains in addition to a backbone of 60 mitochondrial genome sequences published early (Guilliet *et al*. 2022). The phylogenetic reconstruction revealed a clear structuring of the strains into three distinct mitochondrial haplotypes (Figure 2). The Cameroon and Myanmar strains clustered within the haplotype C2 lineage while the Indonesian and European strains grouped within haplotype C1. In contrast, the Chinese strain formed a separate clade corresponding to haplotype D. These lineages were consistent with previously described haplotypic groups reported in worldwide BSF populations with the C1 lineage clustering most of the industrial strains used in Europe. Bootstrap support values indicated strong support for the main nodes of the tree, confirming the robustness of the inferred relationships.

**Figure 2:**
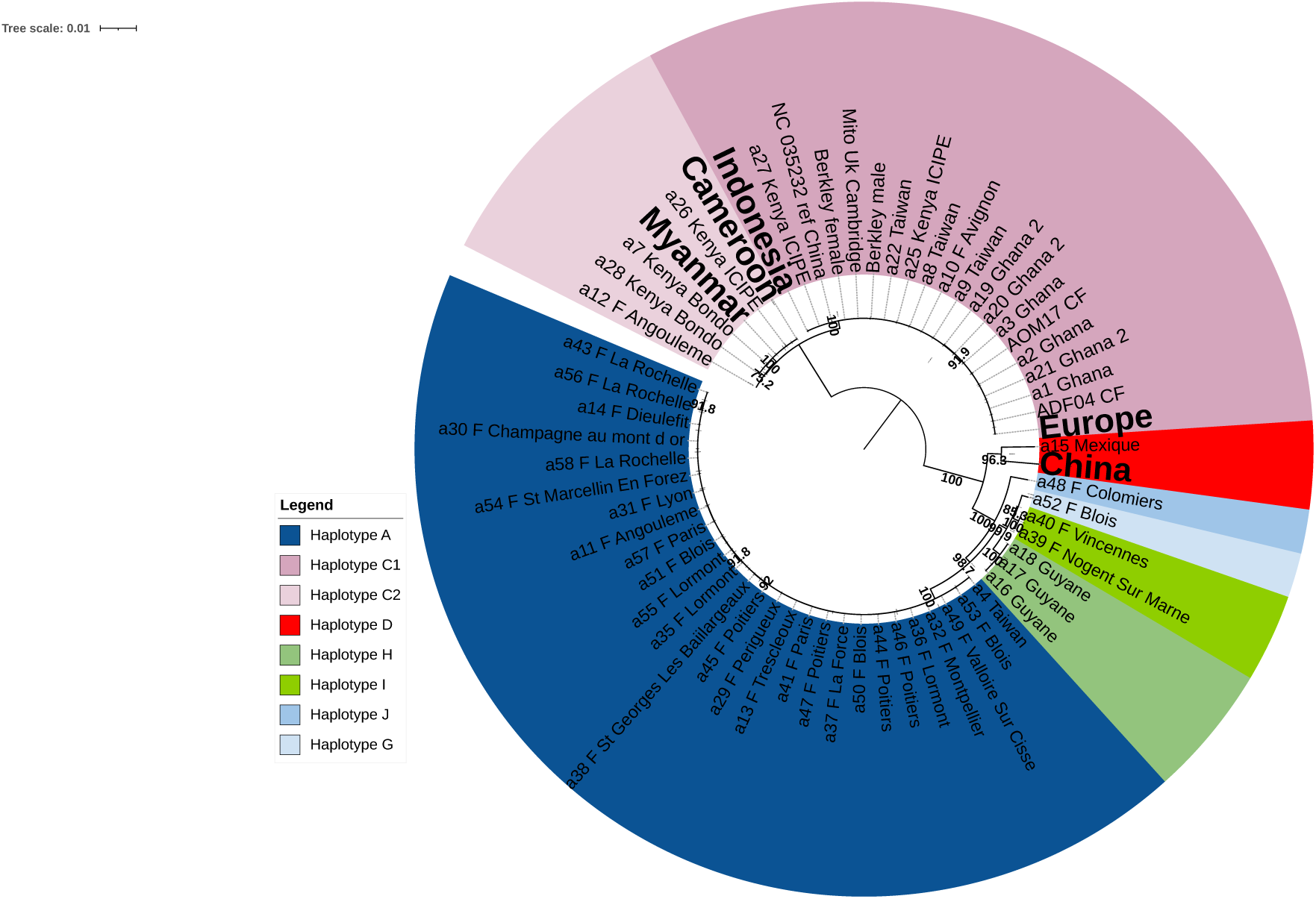
Mitochondrial maximum likelihood phylogenetic tree of the five BSF strains used in this study. Colored segments represent the different mitochondrial haplogroups identified in earlier studies (Guilliet *et al*. 2022). The position of the five strains have been highlighted in bold, bootstrap values are indicated from 75 to 100.

To further investigate the genetic relationships among the five BSF strains, we performed nuclear whole-genome SNP calling. After filtering, we collected a total of 9,495,746 SNPs for the 5 samples. Based on these SNPs, we reconstructed a maximum likelihood phylogenetic tree using nuclear genomic variants that revealed clear genetic structuring among strains (Figure 3A). The European and Burmese strains formed a closely related cluster supported by high bootstrap values, while the Indonesian and Chinese strains grouped together in a separate lineage. The Cameroonian strain appeared genetically distinct. By comparison, the results obtained with nuclear genomic data are very different from those shown on mitochondrial genome data (Figure 3B). Indeed, the position of the Chinese strain is conflictual: in the mitochondrial tree, China strain appears separated whereas it falls as a sister group of the Indonesian strain in the nuclear SNP tree. Similarly, the Cameroonian strain forms a monophyletic group with the Burmese strain in the SNP tree whereas it appeared isolated from the other strains in the mitochondrial phylogeny.

**Figure 3:**
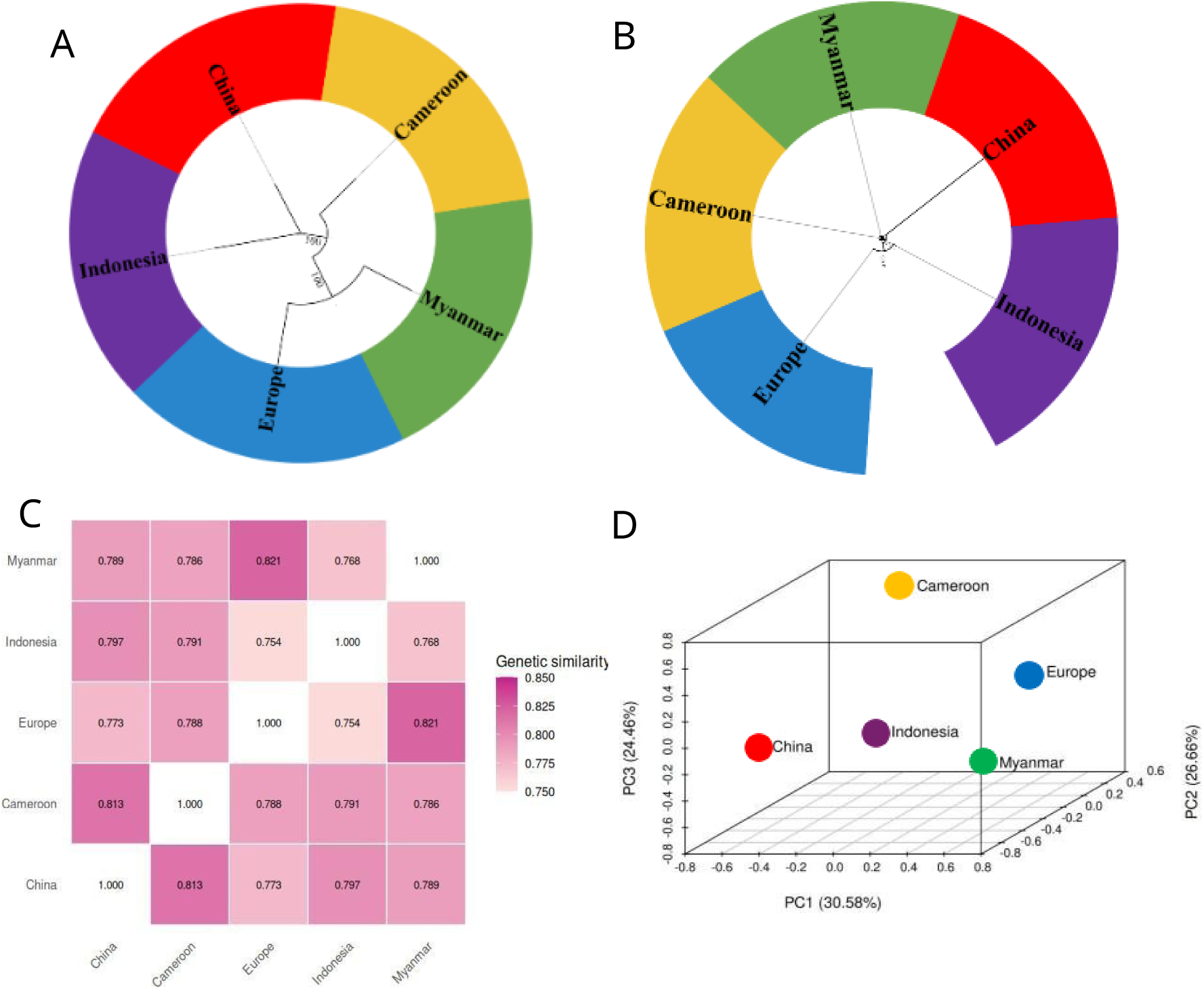
Genetic relationships among the five BSF strains. **(A)** Maximum-likelihood phylogeny inferred from nuclear SNP data illustrating the genetic relationships among the five strains (Cameroon, China, Europe, Indonesia, and Myanmar). (**B)** Maximum-likelihood phylogeny inferred from mitochondrial sequences of the five strains used in the study **(C)** Pairwise genetic similarity matrix among strains based on nuclear SNP variants. **(D)** Principal component analysis (PCA) based on nuclear SNP data, showing the genetic differentiation among strains.

To quantify genetic relationships among strains, a proxy of the pairwise genomic similarity was calculated using SNP sharing and visualized using a heatmap (Figure 3C). Genetic similarity values ranged from 75.4% to 82.1% across strains. The highest similarity was observed between the Europe and Burmese strains (82.1%), whereas lower similarity values were detected between Europe and Indonesia (75.4%), indicating stronger genetic differentiation between these populations in accordance with their respective positions in the nuclear phylogenetic tree.

Population structure was further examined using principal component analysis (PCA) based on nuclear genomic variants (Figure 3D). The first three principal components explained 30.58%, 26.66%, and 24.46% of the total genetic variance, respectively. The PCA separated the five strains in the multivariate space, confirming the genetic differentiation observed in both the nuclear phylogenetic tree and similarity heatmap. In conclusion, our genomic analyses consistently reveal clear genetic differentiation among the five BSF strains, with concordant signals across phylogenetic reconstruction, pairwise similarity, and PCA. However, a notable conflict exists between mitochondrial and nuclear genomic signals, particularly regarding the placement of the Chinese and Cameroonian strains, suggesting incomplete lineage sorting or mitochondrial introgression events. Furthermore, the observed genetic structuring does not follow geographical origin, nor does it reflect the distinction between industrial and artisanal rearing modes, suggesting that strain differentiation reflects independent domestication histories.

### Larval development according to genetic background and diet

Larval growth was monitored over the time for the five strains using three markedly different diets (CF: chicken feed, WB: wheat bran and AM: alfalfa meal). Larval growth trajectories differed markedly depending on both the diet and the strain (Figure 4). Across all strains, larvae reared on chicken feed reached the highest average weights, while individuals reared on alfalfa meal exhibited the lowest growth rates. Wheat bran resulted in intermediate larval weights. Large differences among strains were also observed within each dietary treatment. For instance, the Indonesian and Chinese strains generally exhibited higher larval weights compared to the European strain across most time points. However, the ranking of strains varied depending on the diet, indicating that strain performance was not consistent across substrates. Notably, the Cameroonian strain displayed relatively sustained growth on alfalfa meal compared to other strains, suggesting a greater capacity to utilize nutritionally poor substrates. In contrast, industrially selected strains such as the Indonesian and Chinese strains, despite achieving the highest larval weights on high-quality diets, showed stronger growth depression under low-quality substrate conditions relative to artisanal strains. The European strain consistently underperformed across all dietary treatments, recording the lowest larval weights regardless of substrate quality, indicating a reduced phenotypic plasticity in response to diet variation.

**Figure 4:**
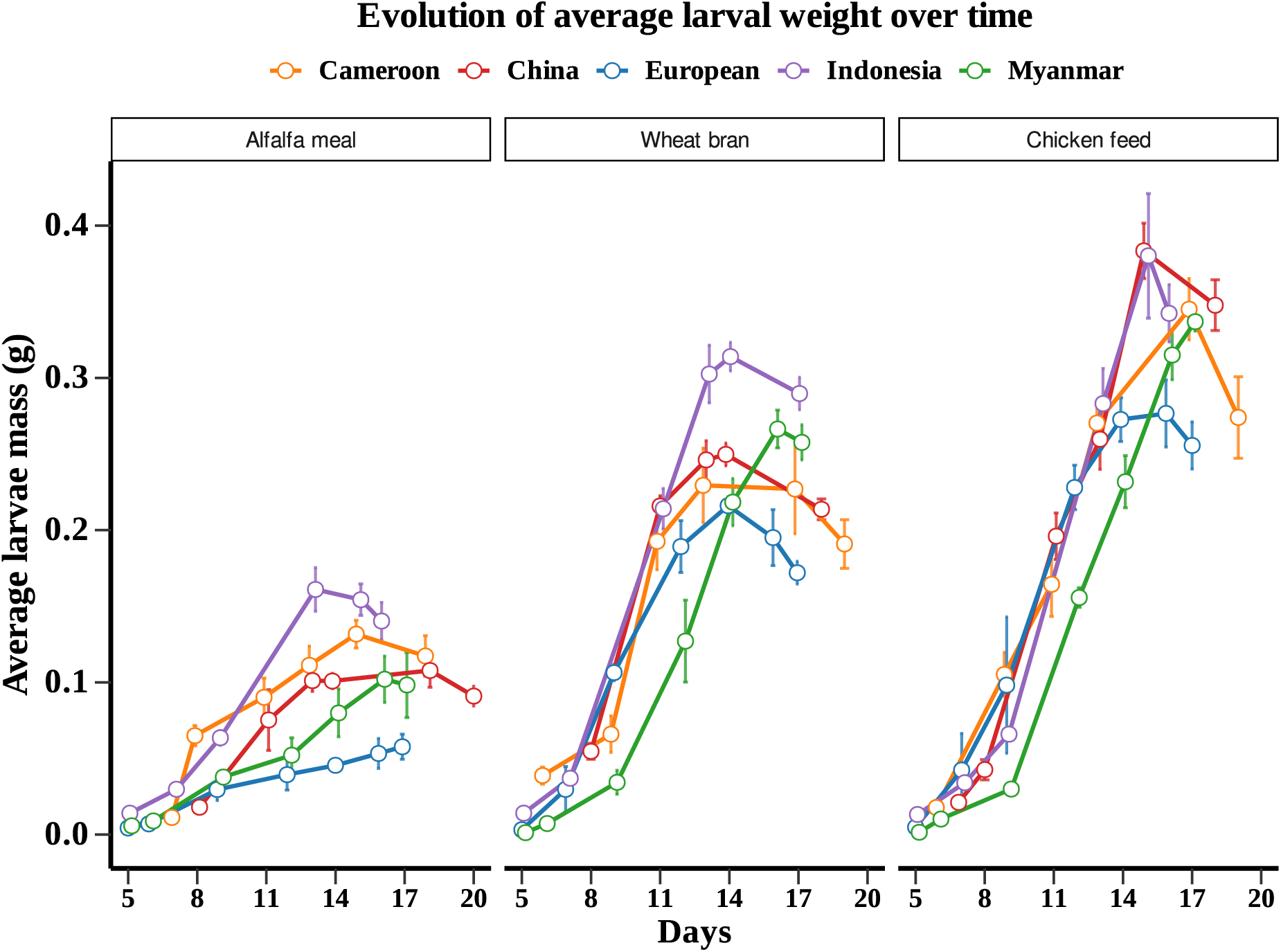
Larval growth dynamics of BSF strains under different diets. Evolution of average larval mass over time for the five BSF strains (Cameroon, China, Europe, Indonesia, and Myanmar) reared on three different diets: alfalfa meal, wheat bran, and chicken feed. Points represent mean larval weight at each sampling time, and error bars indicate standard error using 5 replicates per strain and per diet.

Differences among strains became more pronounced after day 10, when growth trajectories began to diverge substantially. The Indonesian and China strains generally displayed the highest mean weights at later developmental stages, particularly on wheat bran and chicken feed. The Myanmar strain exhibited a slower and more gradual growth pattern across all diets, reaching peak weight later than most other strains. On alfalfa meal, maximum larval weight was reached around day 16–17 across strains, compared to day 14 on wheat bran and chicken feed, indicating that nutritionally poor substrates not only limit growth magnitude but also delay the timing of peak biomass accumulation.

### Interaction of diet and strain on larval life history traits

We compared various life history traits of our five BSF strains according to the three different diets (Figure 5). Detailed quantitative and ANOVA statistical values for all traits across strains and diets are provided in Table 2. The survival rate remained consistently high across all strains and diets, generally exceeding 90%, even for the AM diet that led to small pupae (Figure 5A). No significant differences among diets were detected within strains, indicating that diet composition had limited influence on larval survival under the experimental conditions. This parameter therefore appears relatively conserved across BSF populations.

**Figure 5:**
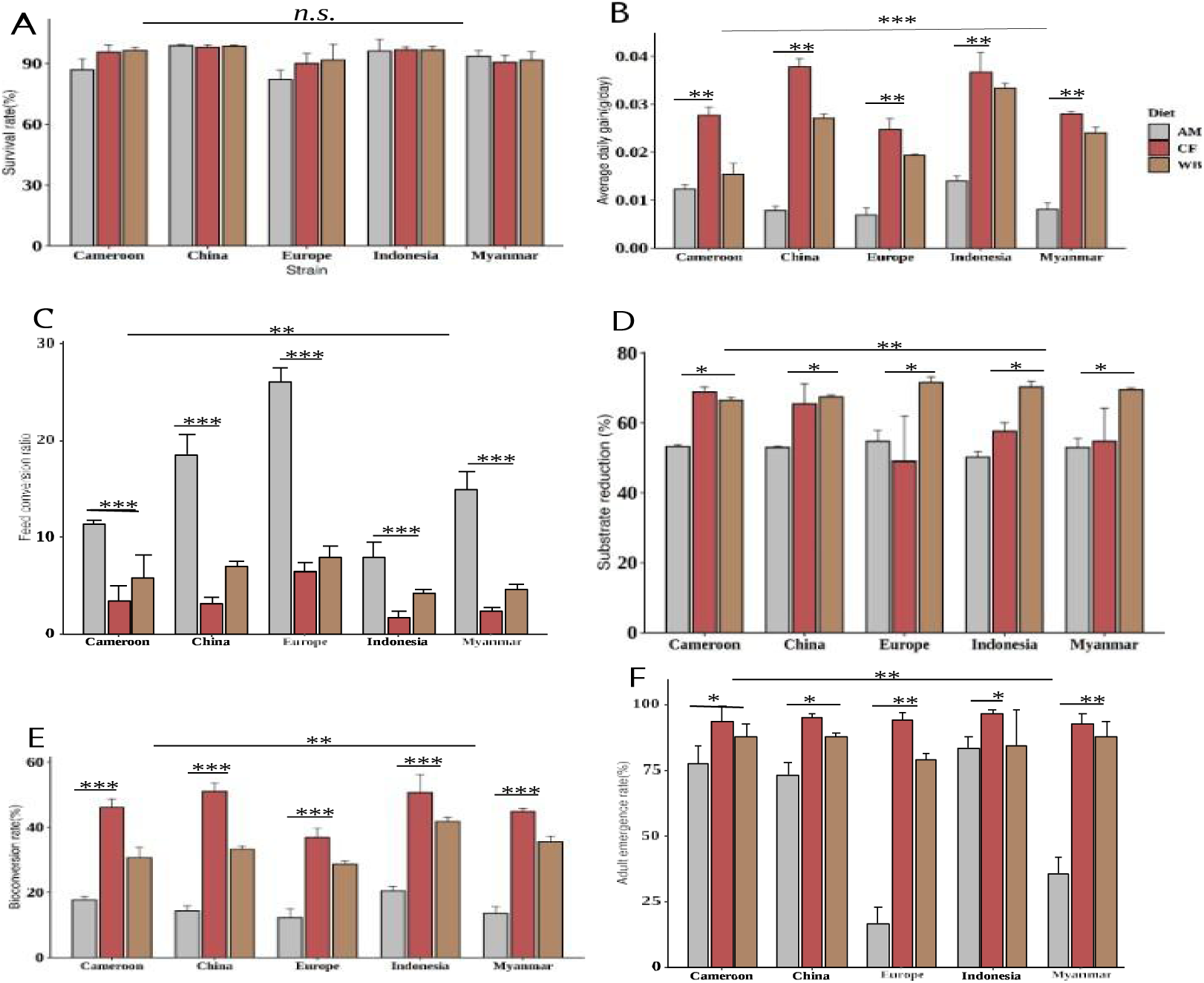
Effects of diet and strain on life-history performance traits of BSF larvae. Larval performance of five BSF strains (Cameroon, China, Europe, Indonesia, and Myanmar) reared on three diets differing in nutritional composition. **(A)** Survival rate (%). **(B)** Average daily gain (g/day). **(C)** Feed conversion ratio (FCR). **(D)** Substrate reduction (%). **(E)** Bioconversion rate (%). (**F**) Adult emergence rate (%). Larvae were reared on three diets: AM (alfalfa meal), CF (chicken feed), and WB (wheat bran). Bars represent mean values ± standard error. Statistical differences among diets within strains are indicated by asterisks using anova test: p < 0.05, ** p < 0.01, *** p < 0.001; **n.s.** indicates non-significant differences.

**Table 2:** ANOVA of the interactions between strains and diets on larval performance and life-history traits in *Hermetia illucens*. Values sharing the same letters indicate that the interactions are not significantly different.

| Strain | Diet | Final weight (g) | Average daily gain (g/day) | FCR | Survival rate (%) | Substrate Reduction (%) | Bioconversion Rate (%) | Adult hatching rate (%) |
| --- | --- | --- | --- | --- | --- | --- | --- | --- |
| China | Alfalfa Meal | 0,108 ± 0,011<sup>h</sup> | 0,008 ± 0,001<sup>fg</sup> | 18,56 ± 2,03<sup>b</sup> | 98,80 ± 0,57<sup>a</sup> | 53,00 ± 0,37<sup>c</sup> | 14,37 ± 1,47<sup>h</sup> | 79,50 ± 6,40<sup>b</sup> |
|  | Chicken Feed | 0,383 ± 0,018<sup>a</sup> | 0,038 ± 0,002<sup>a</sup> | 3,20 ± 0,66<sup>gh</sup> | 98,10 ± 1,08<sup>ab</sup> | 65,44 ± 5,76<sup>ab</sup> | 51,12 ± 2,42<sup>a</sup> | 94,50 ± 2,38<sup>a</sup> |
|  | Wheat Bran | 0,250 ± 0,008<sup>def</sup> | 0,027 ± 0,001<sup>c</sup> | 6,96 ± 0,53<sup>ef</sup> | 98,70 ± 0,45<sup>ab</sup> | 67,47 ± 0,52<sup>a</sup> | 33,29 ± 1,00<sup>de</sup> | 93,50 ± 1,29<sup>a</sup> |
| Cameroon | Alfalfa Meal | 0,132 ± 0,009<sup>gh</sup> | 0,010 ± 0,001<sup>ef</sup> | 11,4 ± 0,37<sup>d</sup> | 86,90 ± 5,45<sup>cd</sup> | 53,27 ± 0,48<sup>c</sup> | 17,56 ± 1,21<sup>gh</sup> | 73,50 ± 11,47<sup>b</sup> |
|  | Chicken Feed | 0,345 ± 0,020<sup>ab</sup> | 0,028 ± 0,002<sup>c</sup> | 3,4 ± 1,56<sup>gh</sup> | 95,50 ± 3,62<sup>abc</sup> | 68,89 ± 1,41<sup>a</sup> | 46,01 ± 2,70<sup>ab</sup> | 91,25 ± 8,66<sup>a</sup> |
|  | Wheat Bran | 0,229 ± 0,024<sup>ef</sup> | 0,028 ± 0,003<sup>c</sup> | 5,76 ± 2,46<sup>ef</sup> | 96,50 ± 1,54<sup>ab</sup> | 66,51 ± 0,81<sup>ab</sup> | 30,59 ± 3,26<sup>ef</sup> | 94,25 ± 1,89<sup>a</sup> |
| Europe | Alfalfa Meal | 0,058 ± 0,008<sup>i</sup> | 0,005 ± 0,001<sup>g</sup> | 26,12 ± 1,4<sup>a</sup> | 82,10 ± 4,68<sup>d</sup> | 54,72 ± 3,18<sup>c</sup> | 7,70 ± 1,10<sup>i</sup> | 2,50 ± 1,00<sup>d</sup> |
|  | Chicken Feed | 0,277 ± 0,022<sup>cd</sup> | 0,025 ± 0,002<sup>c</sup> | 6,46 ± 0,9<sup>ef</sup> | 90,00 ± 5,11<sup>bcd</sup> | 17,83 ± 10,39<sup>d</sup> | 36,88 ± 2,93<sup>cd</sup> | 96,00 ± 4,32<sup>c</sup> |
|  | Wheat Bran | 0,216 ± 0,006<sup>f</sup> | 0,019 ± 0,000<sup>d</sup> | 7,95 ± 1,17<sup>e</sup> | 91,80 ± 7,60<sup>abc</sup> | 71,63 ± 1,51<sup>a</sup> | 28,81 ± 0,78<sup>f</sup> | 92,50 ± 6,19<sup>c</sup> |
| Indonesia | Alfalfa Meal | 0,154 ± 0,010<sup>g</sup> | 0,014 ± 0,001<sup>e</sup> | 7,9 ± 1,58<sup>e</sup> | 96,20 ± 5,72<sup>ab</sup> | 50,23 ± 1,51<sup>c</sup> | 20,57 ± 1,37<sup>g</sup> | 90,25 ± 14,31<sup>a</sup> |
|  | Chicken Feed | 0,380 ± 0,041<sup>a</sup> | 0,037 ± 0,004<sup>ab</sup> | 1,71 ± 0,69<sup>i</sup> | 96,80 ± 1,48<sup>ab</sup> | 57,60 ± 2,45<sup>bc</sup> | 50,68 ± 5,46<sup>a</sup> | 94,75 ± 2,63<sup>a</sup> |
|  | Wheat Bran | 0,314 ± 0,009<sup>bc</sup> | 0,033 ± 0,001<sup>b</sup> | 4,23 ± 0,38<sup>fg</sup> | 96,60 ± 1,92<sup>ab</sup> | 70,31 ± 1,53<sup>a</sup> | 41,85 ± 1,26<sup>bc</sup> | 92,25 ± 7,41<sup>a</sup> |
| Myanmar | Alfalfa Meal | 0,102 ± 0,015<sup>h</sup> | 0,008 ± 0,001<sup>fg</sup> | 14,9 ± 1,9<sup>c</sup> | 93,60 ± 2,86<sup>abc</sup> | 52,97 ± 2,63<sup>c</sup> | 13,61 ± 2,03<sup>h</sup> | 5,50 ± 6,40<sup>d</sup> |
|  | Chicken Feed | 0,337 ± 0,006<sup>b</sup> | 0,028 ± 0,001<sup>c</sup> | 2,4 ± 0,39<sup>hi</sup> | 90,60 ± 3,49<sup>abcd</sup> | 21,02 ± 8,32<sup>d</sup> | 44,92 ± 0,81<sup>b</sup> | 96,50 ± 2,52<sup>c</sup> |
|  | Wheat Bran | 0,266 ± 0,012<sup>de</sup> | 0,024 ± 0,001<sup>c</sup> | 4,63 ± 0,47<sup>fg</sup> | 91,80 ± 4,01<sup>abc</sup> | 69,52 ± 0,53<sup>a</sup> | 35,52 ± 1,65<sup>de</sup> | 96,50 ± 3,00<sup>c</sup> |

By contrast, all the other parameters display strong differences among strains and diets. Average daily gain (ADG) was strongly influenced by diet across all strains (Figure 5B). For each strain, larvae reared on chicken feed (CF) exhibited significantly higher growth rates compared to those fed wheat bran (WB) and alfalfa meal (AM) (p < 0.01, p < 0.001). Conversely, AM consistently resulted in the lowest ADG values. Among strains, the Chinese and Indonesian strains showed the highest growth performance under CF, reaching approximately 0.037–0.038 g/day. In contrast, the European strain exhibited comparatively lower growth across all diets, particularly under AM, where ADG values were minimal (∼0.007 g/day). Wheat bran generally produced intermediate growth rates, although its effect varied depending on the strain, highlighting a genotype-dependent response to diet. Feed conversion ratio (FCR), which measured the efficiency with which larvae convert feed into biomass, was also strongly influenced by diet (Figure 5C). Across all strains, larvae reared on AM exhibited significantly higher FCR values compared to those fed CF or WB (p < 0.01 to p < 0.001), indicating lower feed efficiency on this substrate. Marked differences were also observed among strains. The European strain displayed the highest FCR values across diets, particularly under AM (approximately 26), reflecting poor feed efficiency. In contrast, the Indonesian strain exhibited the lowest FCR values, especially under CF diet (approximately 2), indicating better substrate conversion efficiency. The Chinese and Myanmar strains showed intermediate responses, while the Cameroon strain demonstrated relatively stable but moderate efficiency across diets. Substrate reduction differed significantly among diets (Figure 5D). In most cases, the highest reduction values were observed with CF, followed by WB, whereas AM consistently resulted in lower reduction levels (p < 0.05 for several strains). A similar pattern was observed for bioconversion rate (Figure 5E), which varied significantly among diets (p < 0.001 for most strains). Larvae reared on CF consistently achieved the highest bioconversion values, whereas WB showed intermediate performance and AM resulted in the lowest values. Similarly, adult emergence rate was significantly influenced by diet across all strains (Figure 5F). Larvae reared on CF consistently exhibited the highest emergence rates, generally exceeding 90%, whereas AM resulted in the lowest emergence rates, with a particularly strong reduction observed in the European strain (∼15–20%) (p < 0.05 to p < 0.01) consistent with the markedly lower larval weights recorded for both the European and Myanmar strains on alfalfa meal, suggesting that insufficient larval biomass accumulation under nutritionally poor conditions compromises successful pupation and adult emergence. Wheat bran produced intermediate emergence rates across all strains. Substantial differences were also observed among strains, with the Chinese and Indonesian strains showing consistently high emergence across diets, while the European strain exhibited a strong sensitivity to suboptimal diet conditions.

Finally two-way ANOVA revealed a significant strain × diet interaction for all performance traits except survival rate (Table 2), indicating that the relative performance of strains was not consistent across dietary treatments. This interaction was particularly evident for feed conversion ratio (Table 2) : the European strain showed a disproportionately high FCR on alfalfa meal compared to all other strains, while the Indonesian strain maintained consistently low FCR values regardless of diet. Similarly, bioconversion rate and average daily gain displayed strain-specific responses to diet quality, with the Chinese and Indonesian strains showing the steepest performance decline from chicken feed to alfalfa meal, whereas the Cameroonian strain maintained more stable bioconversion across substrates. Substrate reduction followed a similar pattern, with wheat bran and chicken feed consistently yielding higher reduction rates than alfalfa meal across all strains, though the magnitude of this difference varied among strains. Survival rate was the only trait unaffected by the interaction (n.s.), remaining high and stable across all strain × diet combinations (Table 2), suggesting that larval viability may be buffered against both genetic background and substrate quality under the conditions tested. Overall, although diet represented an important factor influencing larval performance, consistent differences among strains were observed across multiple traits for a given diet. In particular, the Chinese and Indonesian strains generally exhibited higher performance, whereas the European strain tended to display lower growth and conversion efficiency across diets.

## Discussion

Now considered a species of major importance for the feed industry, *Hermetia illucens* has gained significant attention in recent years due to its capacity to valorize a wide range of organic substrates and produce high-value protein and fat. In this context, understanding the genetic diversity of BSF populations becomes essential for optimizing strain selection, improving performance, and ensuring long-term sustainability of breeding programs.

In this work, we have first genetically characterized five strains of BSF collected from artisanal and industrial farms and show that there is a high level of genetic diversity among these BSF strains. Following this, we try to understand if these genetic differences correlated with various life history traits rearing the larvae on three different diets. Our results revealed that diet quality was logically an important driver of larval growth, bioconversion, and feed conversion efficiency but strain effects and strain × diet interactions were also significant for most traits illustrating a complex interplay between the genetic makeup of each strain and the different rearing conditions.

### Genetic structuring among BSF strains

Our genomic analyses revealed a clear genetic differentiation among the five *Hermetia illucens* strains investigated in this study. Both mitochondrial and nuclear genomic approaches consistently indicated that these strains belong to distinct genetic lineages within the global diversity of the species. Similar patterns of genetic structuring have previously been reported in worldwide BSF populations, suggesting that the species is composed of multiple differentiated lineages rather than a single homogeneous population (Stahls *et al*., 2020; Kaya *et al*., 2021). Mitochondrial phylogeny placed the strains into three previously described haplotypic groups, haplotype C1 including Indonesia and Europe strain, haplotype C2 with strain from Burmese and Cameroon (Both this haplotype are commerial one) and haplotype D with strain from China; consistent with the global haplotype structure identified in earlier phylogeographic studies (Guilliet *et al*., 2022). The inclusion of sequences that are accessible to the public confirmed that the strains analyzed in this study are distributed across distinct maternal lineages found in different geographic regions. Such patterns likely reflect the complex evolutionary history of BSF populations, which have been shaped by both natural dispersal and recent human-mediated introductions associated with the rapid expansion of BSF farming (Barragan-Fonseca *et al*., 2017). Interestingly, the mitochondrial and nuclear phylogenies revealed partially different relationships among the strains. While mitochondrial genomes reflect strictly maternal inheritance, nuclear genomes integrate biparental inheritance and recombination, which may produce distinct evolutionary signals. Discordances between mitochondrial and nuclear markers have been widely documented in population genomic studies and may result from historical introgression events or population admixture (Toews and Brelsford, 2012). Limited introgression and gene flow between divergent populations have been recently observed in BSF (Generalovic *et al*., 2023). In the context of BSF domestication, such discrepancies may also reflect recent mixing of strains originating from different breeding programs. These results have to be validated by population genomic analysis based on resequencing data on multiple individuals of each strain. Indeed, another possible explanation lies in the limited number of strains and individuals included in the present analysis. With a small number of sampled populations, phylogenetic reconstruction may be more sensitive to sampling effects, potentially affecting the resolution of certain branches or the inferred relationships among lineages. Increasing the number of strains and representing a broader geographic and genetic diversity could therefore provide a more comprehensive view of the evolutionary relationships among domesticated BSF populations and among wild and captive populations.

Importantly, the genomic analyses indicate that the strains used in this study capture a substantial portion of the genetic diversity observed within the species. Previous studies have emphasized the importance of maintaining genetic diversity in BSF breeding programs, as genetic variation represents the raw material for selection and adaptation to different rearing environments (Barragan-Fonseca *et al*., 2017; Kaya *et al*., 2021). The diversity observed here therefore provides a valuable framework for investigating how genetic background influences phenotypic performance. We therefore investigated whether this genomic differentiation translates into differences in life-history traits when strains are reared under contrasting dietary conditions.

### Diet influences larval performance

In the present study, we confirmed that diet is a major factor influencing larval performance traits in BSF. Larvae reared on chicken feed consistently exhibited higher weight gain, average daily gain, and bioconversion efficiencies compared with those reared on wheat bran or alfalfa meal. These results are consistent with previous studies showing that the nutritional composition of the substrate strongly affects BSF larval growth and biomass production (Nguyen *et al*., 2015; Barragan-Fonseca *et al*., 2017; Sandrock *et al*., 2018, Generalovic *et al*., 2025).

The higher performance observed on chicken feed likely reflects its complex composition with a more balanced nutritional profile, particularly in terms of protein and energy content. In contrast, the lower performance observed on alfalfa meals may be related to reduced digestibility or nutritional limitations of this substrate, as seen on the quantity of frass left. Similar patterns have been reported in several studies demonstrating that substrates with higher protein availability generally support faster larval development and improved biomass conversion efficiency (Gold *et al*., 2018).

Interestingly, differences among diets were also reflected in substrate reduction and bioconversion rates, suggesting that not all substrates are converted into larval biomass with the same efficiency, for instance the Chinese strains achieved a bioconversion rate with around 50% on chicken feed compared to less than 15% on Alfalfa Meal, illustrating how a nutritionally poor substrate can drastically limit the conversion of organic matter into larval biomass. Such differences highlight the importance of substrate quality in determining the efficiency of BSF production systems.

### Genetic background contributes to phenotypic differences

Beyond the strong influence of diet, clear differences in performance were also observed among strains. Across dietary treatments, the Chinese and Indonesian strains generally exhibited higher growth performance, whereas the European strain showed comparatively lower weight gain and growth rates. These results suggest that BSF populations may differ in their capacity to convert organic substrates into biomass. The general poor performance of the European strain compared to the other strains is puzzling. This strain corresponds to the genetic lineage massively used by many companies worldwide. It’s possible that this strain is highly tailored to specific rearing conditions that differ from those used in our experimental setup (higher larval density and/or higher competition for available food). Alternatively, this strain may have undergone strong size population reduction, leading to inbreeding depression and general reduction of its performance as observed in laboratory experiments (Silvaraju *et al*., 2026). It should be valuable in future analysis to compare the diversity of the genetic composition of each population and correlate these parameters with the zootechnic performances obtained in this study.

Genomic analyses conducted in this study revealed substantial genetic differentiation among strains, as evidenced by both mitochondrial and nuclear genome analyses. Similar genetic structuring has been reported in global BSF populations, reflecting the complex evolutionary history and widespread distribution of the species (Stahls *et al*., 2020; Kaya *et al*., 2021). Interestingly, there does not seem to be a link between the strain phylogenetic relatedness based on nuclear data and their respective zootechnic performances. For instance, the Indonesian and European strain cluster together in the nuclear phylogeny yet display markedly contrasted performance profiles across all traits, while Indonesia and Chinese strain grouped in separated clusters but have the highest performances. This leads to thinking about local adaptation among species.

The existence of genetically differentiated lineages suggests that part of the phenotypic variation observed in larval performance traits may be associated with differences in genetic background among strains. Previous studies have suggested that BSF populations maintained in different geographical regions or breeding environments may diverge in their biological performance due to local adaptation or early stages of domestication (Kaya *et al*., 2021). In the present study, the strains originated from both artisanal and industrial breeding contexts, which may also contribute to differences in performance through distinct rearing histories or selection pressures.

Another factor that may contribute to performance variation is the larval microbiota. Several studies have shown that microbial communities associated with BSF larvae can influence digestion efficiency, nutrient assimilation, and growth performance (Bruno et al., 2019) demonstrated that, the gut microbiota composition of BSF larvae differs significantly across gut region and is strongly shaped by diet, with substrate specific bacterial communities likely modulating nutrient breakdown and assimilation capacity. Similarly, Klammsteiner et al. (2020) showed that the core gut microbiome of BSF larvae reared on low-diets remains relatively stable, yet its composition shifts in response to dietary conditions, suggesting that substrate quality directly influences microbial environment available to larvae. As microbiota composition was not characterized in the present study, its potential contribution to the observed inter-strain differences cannot be assessed and represents an important avenue for future research.

### Interaction between diet and genetic background

Importantly, the present results also revealed a significant interaction between diet and strain, indicating that the relative performance of strains varied depending on the substrate. Such genotype × environment interactions are common in biological systems and indicate that the effect of environmental factors cannot be interpreted independently of the genetic background (Hill *et al*., 2004). Similar patterns have been reported in insects, where genetic background modulates the response of individuals to environmental variation (Sandrock *et al*., 2017). In practical terms, this means that a strain performing well on a particular substrate may not necessarily exhibit similar performance on another substrate. These results therefore highlight the importance of considering both diet composition and genetic background when evaluating the production potential of BSF strains. Accounting for genotype × environment interactions may thus be critical for the development of efficient breeding and rearing strategies in BSF production systems.

It should also be noted that the present experiments were conducted over a single generation. Although this design allows the detection of strain-specific responses to different diets, it does not allow assessment of the stability of these genotype × environment interactions across multiple generations. Indeed, progressive adaptation to specific diets after a dozen generations has been observed in the BSF (Gligorescu *et al*., 2023; Generalovic *et al*., 2023) indicating that further multi-generational studies would therefore be valuable to determine whether the observed performance differences persist under long-term rearing conditions.

### Implications for BSF domestication and genetic management

Taken together, these findings have important implications for the domestication and optimization of BSF production systems. While optimizing substrate composition remains essential for improving larval growth and biomass production, the observed differences among strains suggest that genetic diversity found in natural populations eventually improved by artificial selection could also play a significant role in improving production efficiency. To translate these results into practical applications, a decision-oriented framework summarizing optimal strain–diet combinations according to specific production objectives is presented in Table 3. This framework highlights that different performance criteria (biomass yield, feed efficiency, substrate reduction) are maximized by different strains and dietary conditions, emphasizing the need for targeted optimization strategies.

**Table 3:** Recommended strain–diet combinations for optimizing production traits in *H. illucens*.

| Target criterion | Recommended strain | Optimal substrate | Justification |
| --- | --- | --- | --- |
| Maximum biomass yield | China, Indonesia | Rich diet | Highest larval weight (China CF: 0.383 g; Indonesia CF: 0.380 g) |
| FCR | Europe, Myanmar | Rich diet | Most efficient feed/biomass conversion |
| Bioconversion rate (BR) | China, Indonesia | Rich diet | Highest biomass production per substrate invested (China CF BR: 51.12) |
| Substrate reduction (SR) | Indonesia, Cameroon | Average diet<br>Rich diet | Best substrate feed efficiency (Cameroon CF SR: 68.89%. Indonesia WB SR: 70.31%) |
| Substrate versatility | Indonesia | All | High and robust performance on CF, WB, and AM |
| Fibrous substrate utilization | Indonesia | Poor diet | Only strain to show acceptable performance (Weight: 0.154 g; BR: 20.57) |

Maintaining and exploiting genetic diversity among BSF populations may therefore represent a valuable strategy for developing strains adapted to specific rearing conditions or substrates. Integrating both environmental optimization and genetic improvement strategies may ultimately contribute to more efficient and sustainable BSF farming systems.

### Conclusion

This study demonstrates that both diet and genetic background play a key role in shaping larval performance in *Hermetia illucens*. Larval growth, feed conversion ratio, substrate reduction, and bioconversion efficiency were strongly influenced by the nutritional quality of the substrate, with chicken feed supporting the highest performance across strains. In addition, clear differences among strains were observed, consistent with the genetic differentiation revealed by mitochondrial and nuclear genomic analyses. The significant interaction between diet and strain indicates that larval performance depends on the combination of genetic background and nutritional environment. These findings highlight the importance of integrating both substrate optimization and genetic selection into future BSF breeding programs. Our work also pointed out the importance of preserving the natural genetic diversity of the BSF that represents an invaluable genetic resource with wild strains suited to relevant diets or rearing conditions and directly usable by farmers for their specific needs. Such approaches could contribute to improving the efficiency and stability of insect-based bioconversion systems.

## Ethics approval

Not applicable

## Data availability

The raw sequencing data generated in this study have been deposited in the NCBI Sequence Read Archive (SRA) under the NCBI BioProject accession [PRJNA1451792](https://www.ncbi.nlm.nih.gov/bioproject/1451792) with the following accessions: SRR38066191 (Myanmar), SRR38066178 (Europe), SRR38066156 (Cameroon), SRR38066106 (Indonesia) and SRR38066136 (China). Raw data and other supplementary datasets are available through the Zenodo community at: https://zenodo.org/communities/anr_insection/, under the DOIs: 10.5281/zenodo.19497246 for ANOVA tests on larval life history trait statistics and R code and 10.5281/zenodo.19497644 for sequence datasets including tree files and alignments.

## Author Contributions

Conceptualization: ADP, POM and JF; Data curation: JJ-L, MM, MK, CG, MF, JF and POM; Formal analysis: JJ-L, MM, ADP, JF and POM; Funding acquisition: JF and POM; Visualization: JJ-L and MM; Writing - original draft: JJ-L and JF; and Writing - review & editing : All authors.

## Declaration of Generative AI and AI-assisted technologies in the writing process

The authors did not use any artificial intelligence assisted technologies in the data analysis and the writing process except for sentence reformulation and spelling.

## Acknowledgments

We would like to thank Isabelle Clavereau and Pascaline Venon for their technical help, Julien Chuche and Guillaume Baudouin for their advices regarding BSF rearing and Samir Mezdour for his help on grant management.

